# Emergent multidien cycles from partial circadian synchrony

**DOI:** 10.1101/2025.07.03.662941

**Authors:** Marc G. Leguia, Maxime O. Baud, Ralph G. Andrzejak

## Abstract

Over the past decades, chronobiology has attracted great attention thanks to the elucidation of the molecular mechanisms underpinning the circadian cycle. Now, growing evidence suggests that cycles longer than circadian, so-called ‘multidien’ cycles, are of crucial importance in physiological fluctuations spanning multiple days with repercussions in health and disease. Unlike circadian clocks, multidien cycles may not be genetically encoded, given their heterogeneity within and across individuals and systems. Here, we propose that multidien cycles may be generated by the interaction between partially-coupled circadian oscillators. To demonstrate this possibility theoretically, we use a ring model of coupled circadian oscillators and study how synchrony within this network evolves over time. We found that a free-running, about-weekly period robustly emerges from the network’s dynamics. A range of additional multidien cycles resulted from subtle variations in the coupling parameters within the network with periodicities reminiscent of those observed across different species. Thus, our model of emergent multidien cycles from partial circadian synchrony constitutes a credible hypothesis for explaining the timing of a myriad of events on the scale of weeks and months in health and disease.

## I. INTRODUCTION

Life is made of intricate biological cycles at multiple timescales. The field of chronobiology^1^, which studies the temporal structure of life, has made striking advances in elucidating the mechanisms^2,3^ of circadian clocks^4–6^ and their role in timing essential physiological functions in anticipation of daytime and nighttime. Recent evidence has shown that longer cycles co-exist with circadian rhythms, in humans^7–12^ and other animals^13^, yet their underpinnings have barely been investigated.

For example, about-monthly cycles may not be limited to women’s menstrual period, as similar timing also exists in testicular functions and hormones^14^. The term ‘multidien cycles’ was coined across fields of investigation^8,10,11,13^ for quasi-rhythms spanning multiple days with variable but shared peak-periodicity centered around 7-10 days (‘about-weekly’^7^) and/or 20-50 days (‘about-monthly’) within and across individuals. In the absence of readily-identifiable external temporal cues (i.e. a ‘Zeitgeber’), these multidien cycles are likely generated endogenously, forming free-running oscillations in which the period-length varies from one cycle to the next. In general, multidien cycles are weaker than circadian cycles, but their importance has recently been emphasized in human health and disease^9,15,16^.

Through the growing use of wearable and implantable monitoring technology, key insights were gained from physiological recordings in natural conditions over weeks, months, or years. Multidien cycles have now been characterized quantitatively in blood pressure^7^, in the heart beat (smart-watch)^9^, in the brain (chronic EEG)^16^, as well as in human behaviour, measured by smartphones^12^. Highly relevant to medicine, multidien cycles can modulate heart-rate by +/-20 beats per minutes^9^ with a potential impact on the timing of cardiovascular events^15,17^. In the brain, multidien fluctuations in cortical excitability may underlie the periodic recurrence of seizures in epilepsy^18–20^, headaches in migraine^21^, and affective states in bipolar disorders^22^, to cite a few. Solid evidence is found in the field of clinical^8,16,23,24^ and experimental epilepsy^25,26^, where multidien cycles of epileptic brain discharges correlate so strongly with the occurrence of seizures that their monitoring enables the forecast of upcoming seizure risk^18,19^.

Thus, multidien cycles are responsible for timing reproduction, vital physiological functions, and are of central importance in brain disorders. Yet, their underlying mechanism remains unknown. A yet-to-be-discovered independent multidien clock that synchronizes physiological systems over days seems unlikely. A rigid hierarchy would hardly explain the broad range of variable periodicities observed across organs. Instead, we here hypothesize that multidien cycles emerge from coupling relationships among many circadian clocks organized in a complex network. We conjecture that the diversity of observed multidien rhythms could result from subtle modifications in coupling parameters, which we demonstrate using the Kuramoto model of coupled oscillators.^27–29^.

## II. METHODS

### A. Model

In our model, we simulate the interactions among *N* = 50 circadian oscillators (e.g. neurons, cells, organs) that are partially coupled among them. We assume that the essential interaction between individual circadian oscillators can be described using only their phase. For this purpose, we use a known modification of the Kuramoto model^27–30^, in which the phase of each oscillator *ϕ*_*j*_(*t*) is described by:

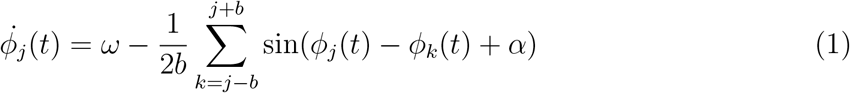

where *ω* is the natural (here circadian) frequency of all coupled oscillators, *b* is the *coupling width*, i.e. how many neighbors on either side each oscillator is coupled to (Fig. 1) and *α* is the *phase lag* parameter. To assess how dynamics may be affected by disrupting the network, we introduce a *coupling reduction* factor *p*, the probability that any link will be removed. An illustration of the network parametrization by *b* and *p* is shown in Fig. 1.

**FIG. 1:**
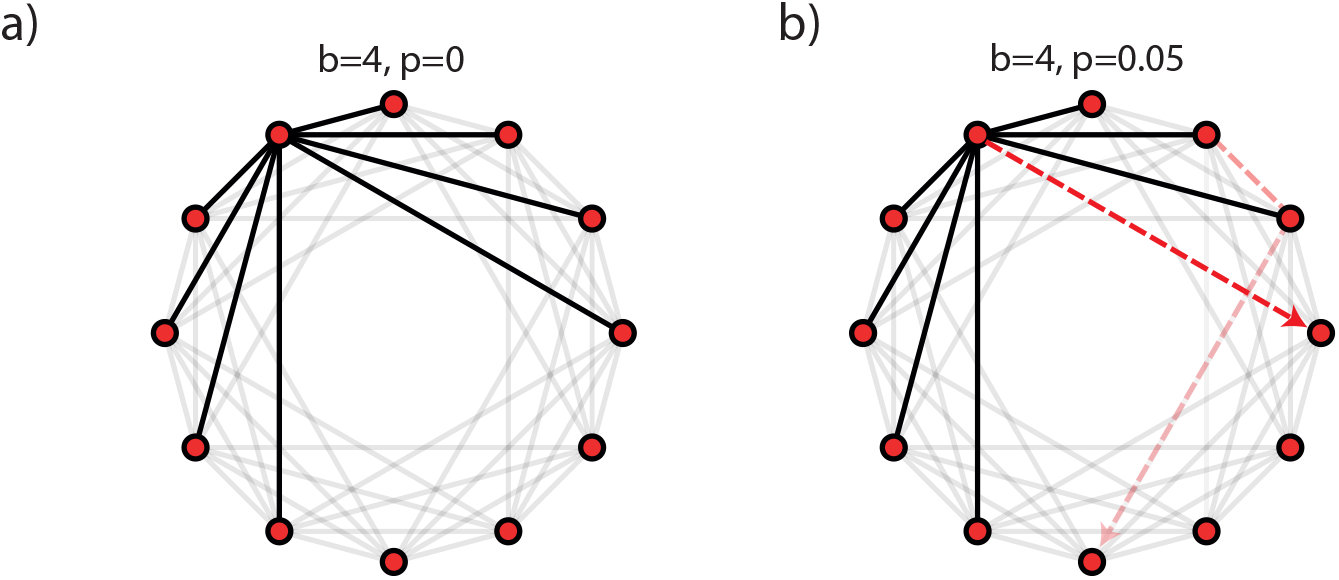
Network of coupled circadian oscillators. a) Illustration of a N=12 network with a coupling width of *b* = 4 and a coupling reduction of *p* = 0. b) Same as a) but with coupling reduction of *p* = 0.05. Here, four links have been removed (red dotted lines). The links with arrow-heads depict links where only one direction has been removed.

For additional analyses, we add an external driver into Eq.1:

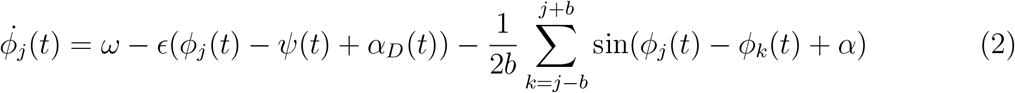

where *ϵ* is the coupling with the external temporal cue *ψ*(*t*), a Zeitgeber with the same natural frequency as the oscillators to mimic synchronization of the circadian rhythm by daylight. Optionally, we included a time-dependent phase lag *α*_*D*_(*t*) with the Zeitgeber, which we kept at *α*_*D*_(*t*) = 0 unless stated otherwise. We set all oscillators to the circadian periodicity *ω*=2*π/*24 and *ϵ* = 0. All simulations of the model are done using a fourth-order Runge-Kutta method with an integration step size of *dt* = 0.05 (72 minutes) and a length of simulations *t* = 5000 days. The overall synchronization of all oscillators is measured by the Kuramoto order parameter^27^, also referred to as the synchronization index^31^, phase-coherence^32^, or phase-locking value^33^ in the literature:

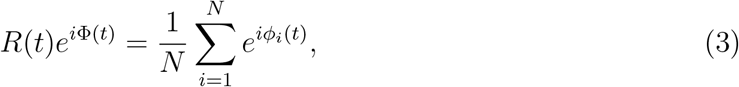

where Φ(*t*) is the average phase and *R*(*t*) is the magnitude of the Kuramoto order parameter, which can take values from *R*(*t*) *→* 0 for a completely asynchronous to *R*(*t*) = 1 for a completely synchronous system of oscillators. As we want to evaluate how *R*(*t*) varies over time, we are only interested in configurations that do not lead to a synchronous state (0 *< R*(*t*) *<* 1) and exclude the realizations that collapse to the fully synchronous state within the simulation time.

Additionally, we model a point-process in which each oscillator ‘spikes’ or ‘releases’ at *ϕ*_*j*_ = 0. The computation of the oscillators’ instantaneous overall spiking allows us to complement the overall synchronization *R*(*t*). Thus, spiking rate *I*(*t*) is computed as the mean of oscillators that crossed the *ϕ* = 0 threshold at each time step of the simulation.

### B. Time-frequency analysis

In our simulation setup, we are interested in how the kuramoto order parameter *R*(*t*) varies over time and whether this variation has some periodicity. To resolve the signal’s instantaneous phase (angle of the Kuramoto order parameter, *e*^*i*Φ(*t*)^) and power (absolute value of the Kuramoto order parameter, *R*(*t*)) at each timepoint on a multidien timescale, we computed a Morlet wavelet transform on *R*(*t*) (or *I*(*t*)) between 4 and 70 days. Using the wavelet transform, we can also compute the periodogram of both *R*(*t*) and *I*(*t*) by averaging the instantaneous power for the simulation time. The first 20000 integration steps (in units of *dt*, corresponding to 1000 simulated days) are discarded to avoid initialization transients.

### C. Surrogate testing

Our null hypothesis *H*_0_ is that the observed periodicity in *R*(*t*) arises from random fluctuations and is not a true feature of the system. To test this, we construct surrogate timeseries by randomly shuffling the original real valued signal *R*(*t*) in time. The surrogate signal *R*(*t*)_*surro*_ retains the original amplitude but has lost all the temporal correlations that may have generated the periodicity we observe. We then use the wavelet transform on the surrogate signal *R*(*t*)_*surro*_ and compare the periodograms of the surrogate signal to the periodogram of the original signal *R*(*t*). We generate 200 of such surrogates, and statistical significance is considered for peaks in the true periodogram that are above the 95^*th*^ percentile of peaks in the surrogate periodograms.

### D. Clustering

Since the system we study has several parameters involved and each set of parameter configurations may lead to different periodicity of both *R*(*t*) and *I*(*t*), we use clustering of the periodograms to extract common patterns across different sets of parameters. Similar to methods in Ref^16^, we used non-negative matrix factorization (NNMF^34^) as an unsupervised clustering algorithm to extract spectral patterns across parameter configurations. Each configuration of parameters *u* is represented by *Q*_*us*_ with *s* corresponding to the indices of the periodogram of length *S* of the Kuramoto order parameter *R*(*t*) calculated with the wavelet transform. We concatenate each configuration periodogram *Q*_*u*_ such as:

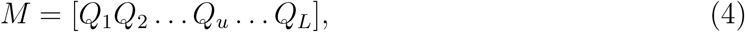

where *L* is the total number of possible configurations of parameters and *M* has *L × S* size. Since matrix *M* does not have negative values, we approximate it (‘factorize’) by two lower-dimensional matrices:

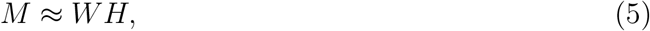

where *W*_*S*_ *× k* and *H*_*k*_ *× L* are the factorized matrices obtained by iterations minimizing the root mean square residual to the original matrix *M* . The matrix *W*_*S*_ *× k* contains the *k* centroids corresponding to the most representative periodograms of length *S*. The corresponding *k* coefficients to approximate each periodogram in M are in matrix *H*_*k*_ *× L*. In our configuration, we use *k* = 5 different clusters to compare them with the clusters found in our previous work^16^.

## III. RESULTS

To obtain a system of *N* = 50 coupled oscillators that are partially synchronized - a so-called chimera state^29,35^, we simulate the model in Eq.1 for coupling widths between *b ∈* [14, 22] oscillators and a phase lag between *α ∈* [1.4, 1.60] radians. For example, in the case of the system of oscillators with *b* = 18 and *α* = 1.46, we observe a dynamic and partial synchronization behavior with some oscillators synchronized and others de-synchronized at any timepoint (Fig 2a). The Kuramoto order parameter oscillates around *R*(*t*) *≈* 0.8 (Fig.2b) without entering full synchrony. We also simulate a point-process *I*(*t*) in which each oscillator ‘spikes’ at a given phase (Fig.2c). Importantly, fluctuations in the overall synchronization (Fig. 2b) and spiking rates (Fig. 2c) showed emergent periodicity centered at *∼* 7 and *∼* 45 days for this set of parameters (Fig.2d).

**FIG. 2:**
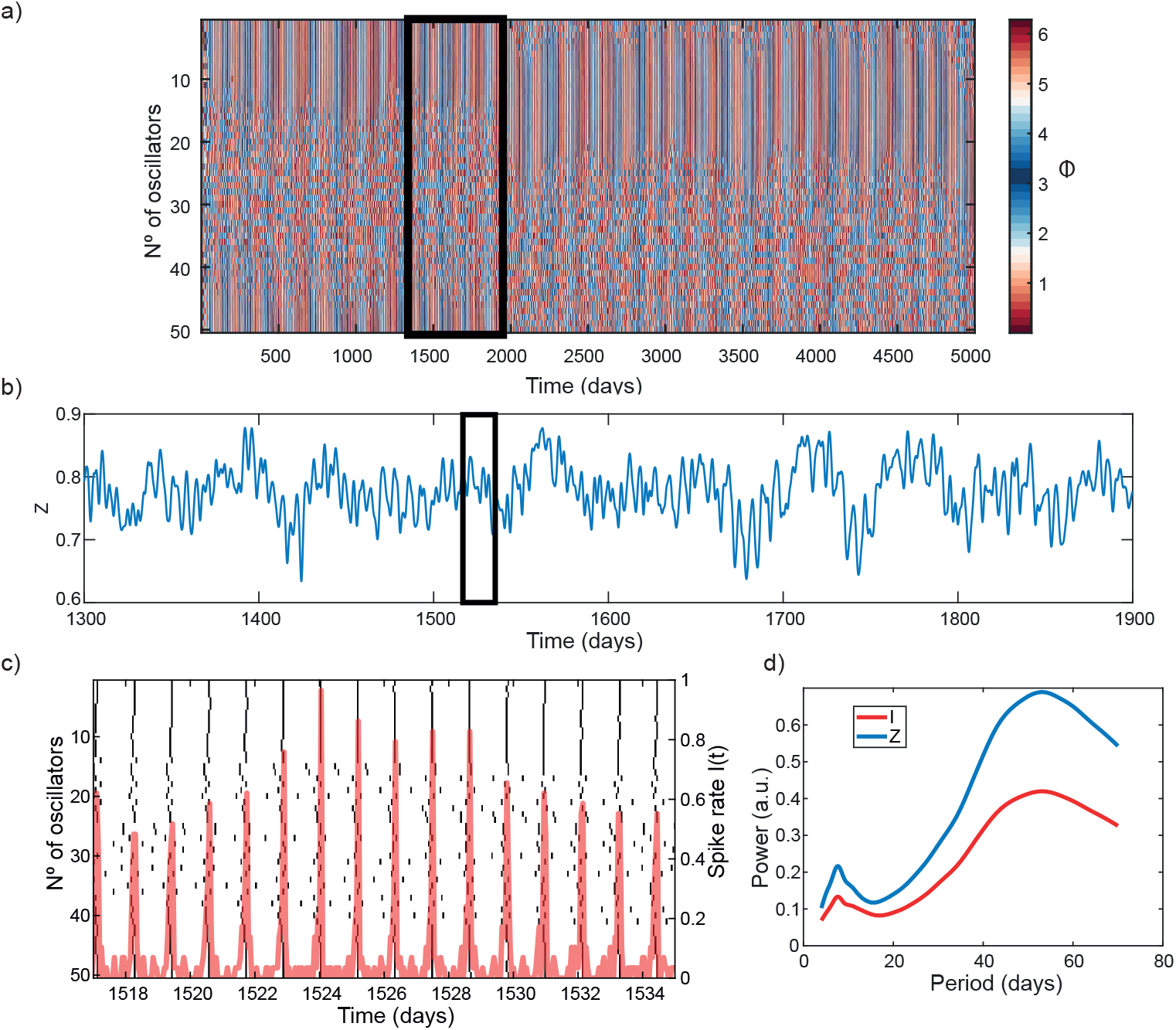
Coupled circadian oscillators generate multidien cycles. a) Partial synchronization formation (so-called ‘Chimera state’) in the interaction of *N* = 50 oscillators with coupling width *b* = 18 and phase lag *α* = 1.46. b) Kuramoto order parameter oscillations over the window depicted in a). c) Discretization of the spiking rates *I*(*t*) across all the oscillators (spiking at *ϕ* = 0) over the window depicted in b). The trace in red shows fluctuations in the spiking rate (y-axis on the right) at circadian (within each day) and multidien (across days) timescales. d) Periodicity of the Kuramoto order parameter *R*(*t*) and the spiking rate *I*(*t*) over the simulation period.

Specifically, the spiking rate *I*(*t*) yields a fast oscillation (circadian) modulated by a slower one (multidien) which resembles periodic fluctuations in inter-ictal epileptiform activity (IEA) recorded with chronic EEG devices analyzed by us^8,16^ and others^8,9,16,23^ in patients with focal epilepsy. Intuitively, firing neurons that lack complete circadian synchronization self-generate slower modulation of their concerted firing. This result from our model indicates that a system of coupled circadian oscillators, each spontaneously oscillating with identical periodicity but at a different phase, is sufficient to generate multidien cycles. As the order parameter *R*(*t*) and spikes *I*(*t*) share the same periodicities, subsequent analyses focused on *R*(*t*).

To characterize periodic fluctuations in the Kuramoto order parameter *R*(*t*) for a range of parameters, we average 10 simulations of the system in Eq.1 for pairs of coupling widths *b* and phase lags *α*. By fixing *α* = 1.52 radians, and varying the coupling width *b*, we observe systematic changes in the periodicity of *R*(*t*). In particular, the broader the coupling width, the longer the multidien periods (Fig.3a). Inversely, by fixing the coupling width at *b* = 15 oscillators and varying the phase lag, we observe varying multidien cycle strength in *R*(*t*) (Fig.3b) with phase-lags close to *π/*2 (anti-phase) intuitively abating multidien periodicity, as partial synchronization is known to disappear for *α > π/*2^35,36^. In general, the maintenance of partial synchronization also depends on consistent symmetric coupling^37,38^. We randomly disrupted coupling within the network of oscillators and evaluated the impact on the periodicity of the Kuramoto order parameter *R*(*t*) (Fig.3c). For a fixed coupling width *b* and phase lags *α*, the periodogram is stable for a low coupling reduction *p*. It loses the longer periodicity (45 days in the example) when the coupling reduction is *p* = 0.05 (Fig.3d).

**FIG. 3:**
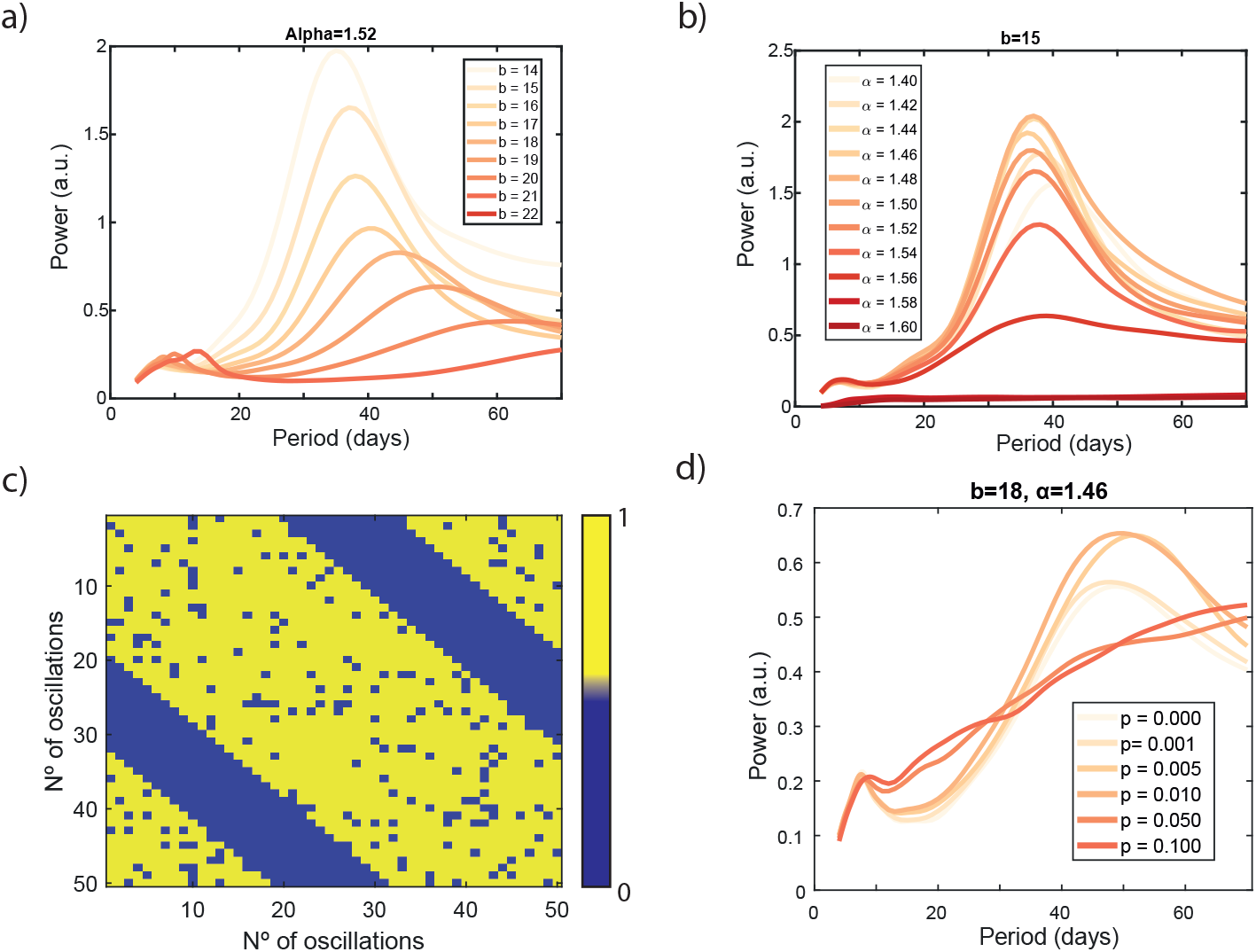
Modulation of multidien cycles by varying circadian coupling. The coupling width *b*, the phase-lag *α*, and coupling reduction *p* contribute differently to the period and power in *R*(*t*). a) Periodogram of the Kuramoto order parameter *R*(*t*) as a function of the coupling width *b* keeping the phase lag parameter fixed at *α* = 1.52. b) Same as a) but fixing *b* = 15 and changing the phase lag parameter *α*. c) Example of the connectivity matrix with randomly removed links for a coupling reduction probability of *p* = 0.1. d) Same as a) with *α* = 1.42 and *b* = 18 with different coupling reduction probabilities *p*.

In the parameter space of our model and for all different configurations of the system changing *α, b*, and *p* we systematically observed the emergence of a *∼* 7 days peak-periodicity in agreement with Halberg’s free-running circaseptan rhythm^39^ and other evidence^9,16^(Fig.4a). Furthermore, using the NNMF unsupervised clustering algorithm, we extract five centroid periodograms across all parameters studied and average periodograms belonging to the same cluster (Fig.4b). While comparing real data directly to a parameter-space exploration must be done with caution, the average periodicities intriguingly, resembled those observed in real brain recordings^16^(Fig.4b). Thus, we find that interactions of circadian oscillators can produce known patterns observed in longer periodicities (multidien) with an attractor periodicity of *∼* 7 days.

**FIG. 4:**
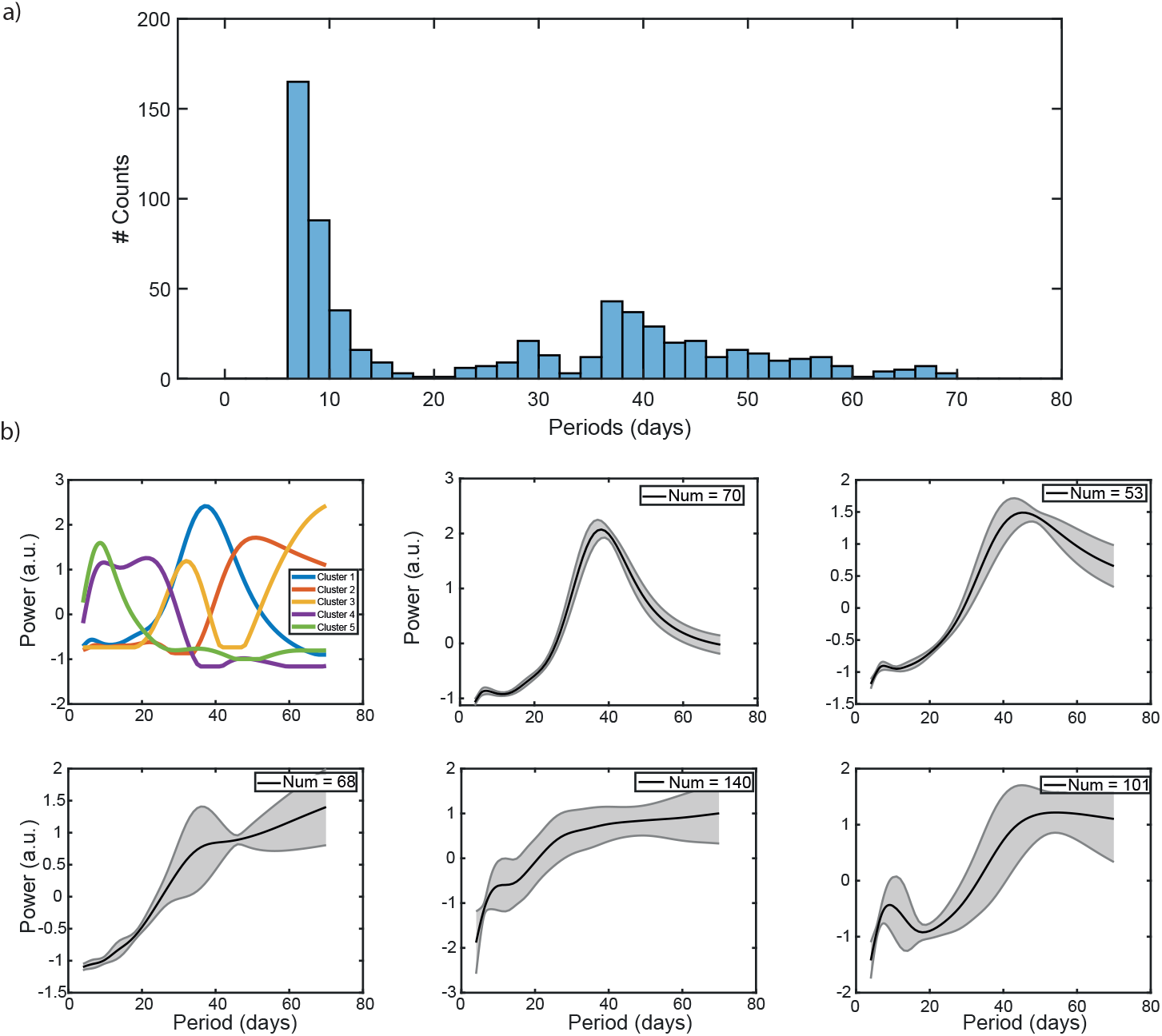
Multidien periodicities found across the paramater space. a) Histogram for all significant peaks (assessed by testing against 200 surrogate timeseries, methods) calculated for all combinations varying *α, b*, and *p*. b) Five clusters were extracted using NNMF. The average periodograms for all the combinations of variables in each of the five clusters are also shown in black with their standard deviation as the grey area.

So far, we simulated the system of Eq.1 in the absence of a Zeitgeber, i.e., keeping *ϵ* = 0, which we now introduce to all oscillators at *t* = 500 days to characterize its impact on multidien cycles, while keeping other parameters fixed. Immediately upon introduction of a circadian Zeitgeber, multidien cycles become stronger Fig.5a. In particular, we observe that the stronger the Zeitgeber, the stronger the *∼* 7 day multidien cycle (lower plot in Fig.5b). This suggests that the environment can have an impact on self-organized multidien cycles, but yet do not generate them.

**FIG. 5:**
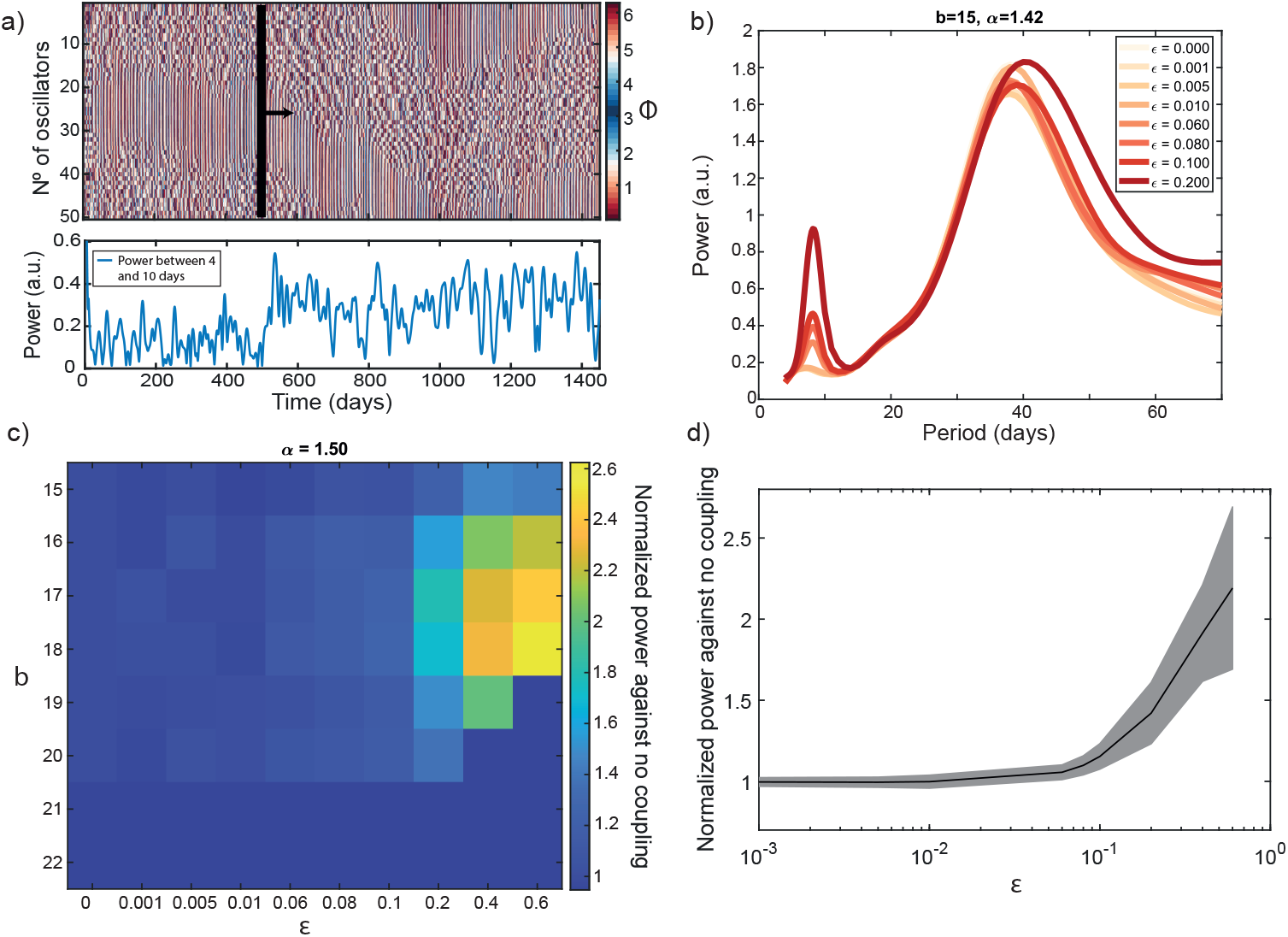
The addition of a circadian Zeitgeber in the system of oscillators enhances the magnitude of multidien cycles. a) Partial synchronization example with an arrow showing when the Zeitgeber is added to the system of oscillators (*ϵ* = 0.1). The lower plot depicts the overall power between 4-10 days of the order parameter oscillations. b) Periodogram of the order parameter as a function of Zeitgeber strength *ϵ*, keeping *b* = 15, and *α* = 1.42. c) Change of the power in the multidien range keeping *α* = 1.50 and varying the coupling width *b* and the Zeitgeber strength *ϵ*. d) Overall power in the multidien range as a function of *ϵ* averaging for the different *α*’s and *b*’s.

## IV. DISCUSSION

Here, we demonstrated how multidien cycles may emerge from the interactions among a network of circadian oscillators. We propose that the network’s variable coupling width (*b*), coupling reduction (*p*) or phase lag (*α*) suffice to give rise to a range of multidien periodicities reminiscent of those observed across health^7,39,40^ and disease^8,9,23^ and in different species^16,25,26^. This model of emerging, self-organized multidien cycles represents a credible alternative to that of a separate master multidien clock that would entrain physiological systems throughout the body. Indeed, it reconciles with the intuition that variable multidien cycles may not be genetically-encoded. Furthermore, characteristics of our network (e.g. Kuramoto order parameter) align with those found in circadian gene expression, showing partial synchronization across 46 human tissues^41^.

As a complex network of many circadian oscillators enter into a state in which some oscillators are synchronized whereas others are not, momentary dynamic interactions generate fluctuations whose period length varies from cycle to cycle. Resulting from this ‘chimera state’, multidien cycles are quasi-rhythms that are non-stationary but revolve around a central period, set by the individual tuning of network parameters. The resulting dynamics can fully account for the variability in multidien periods observed in real data within and across individuals, without the need to invoke genetic factors, lifestyle, the environment, or drug intake.

Remarkably, an about-weekly period (circaseptan) was the multidien period found most consistently across the parameter space tested, even in the absence of an environmental Zeitgeber. Many earlier studies sought explanations for about-weekly rhythms in the rigid organization of the social week^42^. In contrast, our model suggests that physiological about-weekly rhythms may be endogenously generated and may have led humans to adopt a calendar week of seven days. Second in order of prevalence, about-monthly cycles were the strongest. Our observation of their endogenous emergence helps settle the long-standing debate about lunar influences on physiology in terrestrial animals^43^. We also verified that multidien periodicity was maintained in a cycling environment, and found that a 24h Zeitgeber would strengthen multidien cycles, if anything. Thus, a prediction testable in experiments is the expected strengthening of multidien cycles, as a subject is moved from a constant to a cycling environment.

Limitations of this study relate to the assumptions made in our simplified model to address essential dynamics among 50 simulated oscillators (organ-level). In reality, myriads of cellular clocks do not necessarily have the same periodicity as Zeitgebers and may respect a hierarchy of coupling strengths, which may be the focus of future, more complex models. Our foundational study leaves many open questions: could inter-individual variability in multidien cycles result from such subtle parameter changes, with important consequences for rhythmicity in the system as a whole? Even within an individual, could different systems or organs have different multidien cycles, due to such differences? Could multidien cycles change over time due to illness, age, or environmental factors?

In conclusion, our proposed model represents a mechanistic framework to address the complexities of increasingly observed multidien cycles in health and disease. In the Era of long-term physiological monitoring, it is crucial that hypotheses are well formulated and costly experimental interventions at multidien timescale are soundly motivated. The successful management of chronic dynamical disorders in the fields of cardiology, psychiatry, or neurology may depend on the in-depth understanding of complex dynamics arising from biological rhythms.

## V. DATA AVAILABILITY

The codes and data supporting this manuscript’s findings are available at GitHub.

