## Supplementary materials for "Emergent multidien cycles from partial circadian synchrony"

### Supplementary Emerging of multidien-like rhythms using partial synchronization of phase oscillators

In this section, we provide some supportive evidence and supplementary analyses.

#### I. TOTAL NUMBER OF SPIKES LOCK WITH THE DRIVER

Here we explore the effect of the phase lag of the driving $\alpha_D$ on the overall locking between the sum of spikes $I(t)$ and the time. To do that, we count the value of the spiking data $I(t)$ and lock it in the 24-hour time that can be viewed as a phase. More specifically, we divide the 24 hour day in bins and sum all the I(t) that the signal has for that bin hour. Accordingly, we can calculate the phase-locking value between the sum of spikes $I(t)$ and the hourly phase in the same way as the Kuramoto order parameter. When the coupling between the driver and the system is low, the phase-locking value is close to zero, and there is a uniform distribution of the $I(t)$ in the hourly phases. However, as the coupling gets bigger the distribution is more concentrated around a given hour, and the values of the phase-locking value increase. Thus there are some preferred hours when the $I(t)$ is bigger (Supplementary Fig.1a). Additionally, by changing the phase lag of the driving $\alpha_D$ we can change the phase of the locking (similar to the "jet lag" we experiment when we change the phases in which the sun is driving us). Moreover, this is also the case if we change $\alpha_D$ at each time step in a periodic motion during 365 days between $\alpha_D = 0$ and $\alpha_D = \pi/6$. We observe that as the $\alpha_D$ changes, the phase of the looking of the $I(t)$ and the time also changes progressively (Supplementary Fig.1b). This is interesting as some experiments have shown that the change of light input as the seasons change may also lead to different locking between seizures and the time of the hour they appear. In our case, we find a similar effect controlled by the phase lag of the driver $\alpha_D$.

#### II. SMALL VARIATION ON CIRCADIAN CLOCK NATURAL FREQUENCIES

The circadian clock for humans is not exactly 24 hours and there is heterogeneity in which the value can be a bit higher or lower or change due to age[1,2]. We tested our model for small variations of the natural frequency $\omega_0$ and we found that the profiles of periodicities are very similar to the ones found in by keeping $\omega_0 = 24h$ (Supplementary Fig.2a).

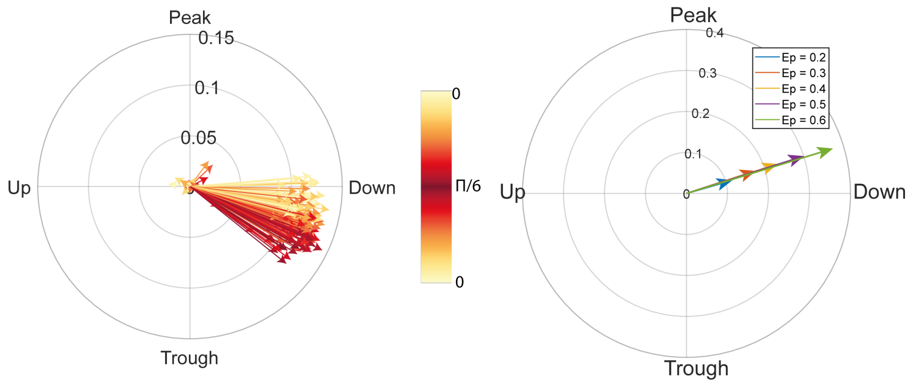


Supplementary Figure 1: **The IEA locking with the sun has a similar effect as the change of locking of $I(t)$ with the time as $\alpha_D(t)$ changes.** a) Locking between the simulated IEA and the time of the day while changing $\alpha_D$ between 0 and $\pi/6$ in one year of simulation. Each arrow represents a monthly average locking which corresponds to 600 datapoints. b) Locking between the IEA and the time of the day for different $\epsilon$ keeping $\alpha_D = -\pi/6$ .

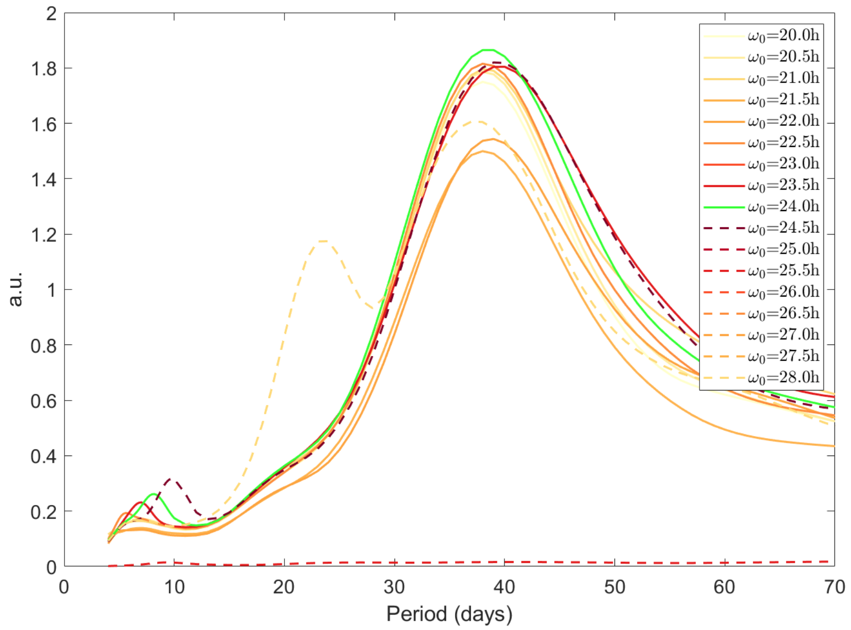


Supplementary Figure 2: **Changing the natural frequency around the 24h period also generates multidien periodicities** a) Periodogram of the model with driving using $\epsilon = 0.05$, $b = 15$ and $\alpha = 1.42$ as a function of the natural frequency $\omega_0$. The green line depicts our original setup with the natural frequency equal to the driver frequency. Solid (dashed) depicts periods lower (higher) than the natural frequency.

### REFERENCES


[1]L. Lack, M. Bailey, N. Lovato, and H. Wright, "Chronotype differences in circadian rhythms of temperature, melatonin, and sleepiness as measured in a modified constant routine pro-

tocol," *Nature and Science of Sleep*, vol. 1, pp. 1–8, Nov. 2009.

[2] J. F. Duffy and C. A. Czeisler, "Age-related change in the relationship between circadian period, circadian phase, and diurnal preference in humans," *Neuroscience Letters*, vol. 318, pp. 117–120, Feb. 2002.
